# Division of Labor between SOS and PafBC in mycobacterial DNA Repair and Mutagenesis

**DOI:** 10.1101/2021.08.05.455301

**Authors:** Oyindamola O. Adefisayo, Pierre Dupuy, James M. Bean, Michael S. Glickman

## Abstract

DNA repair systems allow microbes to survive in diverse environments that compromise chromosomal integrity. Pathogens such as *M. tuberculosis* must contend with the genotoxic host environment, which generates the mutations that underlie antibiotic resistance. Mycobacteria encode the widely distributed SOS pathway, governed by the LexA repressor, but also encode PafBC, a positive regulator of the transcriptional DNA damage response (DDR). Although the transcriptional outputs of these systems have been characterized, their full functional division of labor in survival and mutagenesis is unknown. Here we specifically ablate the PafBC or SOS pathways, alone and in combination, and test their relative contributions to repair. We find that SOS and PafBC have both distinct and overlapping roles that depend on the type of DNA damage. Most notably, we find that quinolone antibiotics and replication fork perturbation are inducers of the PafBC pathway, and that chromosomal mutagenesis is codependent on PafBC and SOS, through shared regulation of the DnaE2/ImuA/B mutasome. These studies define the complex transcriptional regulatory network of the DDR in mycobacteria and provide new insight into the regulatory mechanisms controlling the genesis of antibiotic resistance in *M. tuberculosis*.

## Introduction

Cells are exposed to a variety of endogenous and exogenous factors that lead to DNA damage. DNA damage is particularly relevant to an intracellular pathogen such as *Mycobacterium tuberculosis* as its DNA is subject to assault from a variety of host defense mechanisms, many of which are genotoxic (1-5). There are multiple types of damage that occur on DNA, the most deleterious being double strand breaks. As DNA integrity is essential for cell survival, growth and replication, cells encode multiple genes necessary for DNA damage repair and response (6). The development of antibiotic resistance in *Mycobacterium tuberculosis* is solely the result of chromosomal mutations which arise during replication or because of DNA damage repair and response. A major regulator of the DNA damage response (DDR) in many bacteria is the inducible SOS pathway which is activated when the LexA repressor interacts with RecA nucleoprotein filaments on single strand DNA and undergoes autocatalytic cleavage (7-9). *Mycobacterium* also encodes a second DDR pathway, the PafBC pathway, which functions as a transcriptional activator with a currently unidentified activating signal (10). Although PafBC is encoded in an operon with the ubiquitin-like pup ligase PafA, PafB and PafC do not function in pup-proteasome system (11). Transcriptional analysis of both *M. smegmatis* and *M. tuberculosis* lacking the PafBC pathway (Δ*pafBC*) or *recA* (Δ*recA*, as a surrogate for SOS inactivation) after DNA damage have defined the transcriptional regulon controlled by each pathway (10,12,13). These studies revealed that while there are some important overlaps in the genes regulated by these pathways, the PafBC pathway controls the larger transcriptional output post DNA damage (10,13). The PafBC proteins are not DNA damage inducible and therefore its mechanism of activation is unknown.

However, there are indications that despite having the smaller transcriptional regulon, the SOS pathway has an important functional role. For instance, the SOS pathway has been implicated in the survival of persistent cells (14) as well as in the induction of adaptive mutagenesis (15) in mycobacteria. The DnaE2 polymerase, which is required for mutagenesis in *M. tuberculosis* and *M. smegmatis*, is reported to be under SOS control (15). However, prior literature has used inactivation of RecA as a surrogate for SOS inactivation, due to the essential role of RecA as the LexA coprotease. Although RecA null cells are clearly SOS null, RecA has pleiotropic roles in DSB repair, replication fork restart, and other functions (8,16), raising the possibility that RecA inactivation may have broader effects than simply SOS inactivation (17-19). Our knowledge of the functional overlap between the SOS and PafBC pathways in the mycobacterial DDR is therefore very limited.

This work aims to elucidate how SOS and PafBC functionally contribute to the DNA damage response after distinct types of DNA damage by specifically ablating SOS and PafBC, alone or in combination, in comparison to loss of RecA. To specifically ablate SOS and avoid confounding functions of RecA, we engineered LexA-S167A, which prevents LexA autocatalytic cleavage. Characterization of these bacterial mutants revealed that, although PafBC does control a larger gene set, the two systems are both required for repair, with SOS playing a dominant role after UV damage and PafBC more important for survival after gyrase inhibition.

Studies with specific DNA damaging agents further confirmed that gyrase inhibition and replication fork perturbation specifically activate the PafBC pathway. Finally, we show that, although DnaE2, the primary mutagenic polymerase of mycobacteria, is under SOS control after UV damage, it is coregulated by SOS and PafBC after gyrase inhibition. Further the ImuA/B cassette, also required for mutagenesis, is under dual control and, accordingly, mutagenesis is coregulated by both pathways.

## Results

### Genetic ablation of the DNA damage response in *M. smegmatis*

To investigate the division of labor between the PafBC and SOS DDR pathways, we constructed mutants of each pathway, alone and in combination, in *M. smegmatis*. The PafBC pathway was ablated by deletion of the coding sequences of both proteins (Δ*pafBC*) using a previously validated homologous recombination based knockout strategy (20). LexA is a repressor that is inactivated by RecA stimulated autocatalytic proteolysis (21). As such, deletion of *lexA* results in derepression of SOS, whereas deletion of *recA* may be a poor surrogate for SOS ablation due to its pleiotropic roles in DNA repair. To circumvent this problem, we introduced a point mutation into the *lexA* chromosomal locus to direct synthesis of LexA-S167A, which renders LexA uncleavable (22), in both the wild type (mc^2^155) and Δ*pafBC* strains. These strains (WT, Δ*pafBC, lexAS167A*,Δ*pafBC/lexAS167A*, Δ*recA*), were analyzed by PCR to confirm their genotypes (Fig 1A).

**Figure 1.**
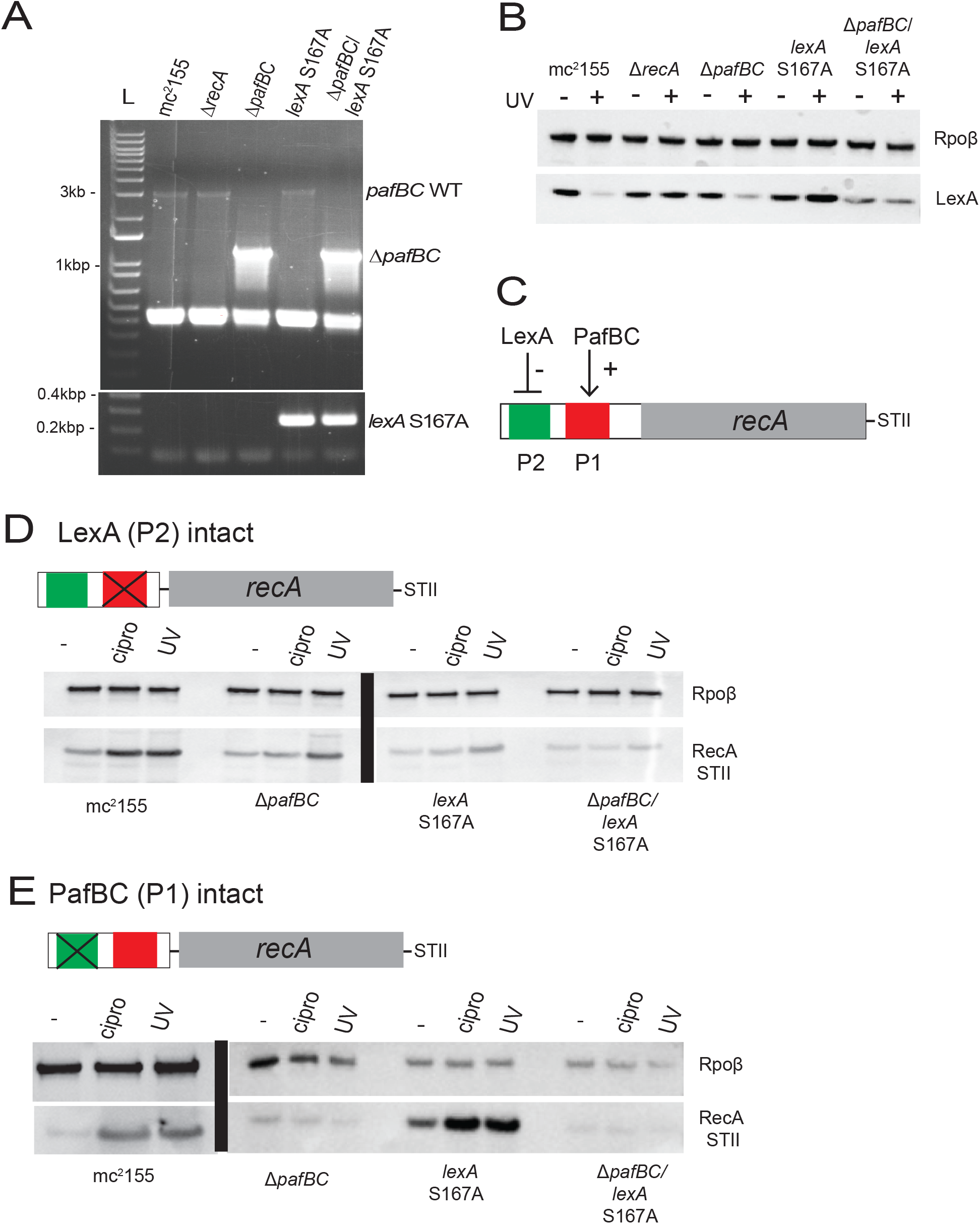
Genetic ablation of the PafBC and SOS pathways in *M. smegmatis*. **(A)** Confirmation of Δ*pafBC* and *lexA*-S167A genotypes of mc^2^155 (WT), Δ*recA*, ΔpafBC, *lexA*-S167A and Δ*pafBC*/*lexA*-S167A strains. The *pafBC* deletion allele was detected by PCR amplification with primers (OAM229 & OAM232) that amplify the genomic region upstream of *pafBC* yielding a product of 3059bp in strains carrying wild type *pafBC* and 1122bp in strains with Δ*pafBC*. The *lexA*-S167A mutation was detected by selective amplification using primers (OAM189 & OAM268) that anneal to the mutated LexA, yielding a product of 234bp for *lexA*-S167A but no product from the wild type allele. L denotes the DNA ladder **(B)** LexA cleavage with DNA damage according to strain genotype. α−LexA immunoblot of mid-log phase expression of LexA (28kD, bottom panel) in mc^2^155, Δ*recA*, Δ*pafBC, lexA*-S167A and Δ*pafBC*/*lexA*-S167A strains without (-) or with (+) DNA damage (20mJ/cm^2^ UV). RpoB is shown as a loading control (top panel). **(C)** Schematic of *recA*-streptag (STII) expression construct containing the PafBC (P1) and LexA (P2) binding sites in the RecA promoter. recA promoter activity was measured by α-streptag (STII) western blot of mid-log phase expression of RecA-STII (37kD) in mc^2^155, Δ*pafBC, lexA*-S167A and, Δ*pafBC*/*lexA*-S167A strains carrying RecA-STII driven only by the LexA repressed P2 (P1 mutated, **D)** or only the *pafBC* promoter (P2 mutated, **E**) with or without DNA damage (UV 20mJ/cm^2^ or ciprofloxacin (1.25μg/ml). RpoB is shown as a loading control for both blots in (D) and (E)

To confirm that the LexAS167A mutation impairs LexA cleavage in vivo, as it did in vitro (data not shown), we analyzed LexA levels by immunoblotting, with or without 20mJ/cm^2^ UV light (Fig 1B). In wild type *M. smegmatis*, LexA protein was detectable at its predicted MW of 28kDa in basal conditions (Fig 1B). With UV treatment, full length LexA became nearly undetectable, although we were unable to detect LexA proteolytic fragments (Fig 1B). In the predicted SOS deficient strains (Δ*recA, lexAS167A* and *pafBC/lexAS167A*), there was no discernible loss of LexA protein with DNA damage (Fig 1B), consistent with impaired LexA cleavage. Ablation of *pafBC* had no effect on LexA cleavage with damage.

To confirm that both pathways, PafBC and SOS, are functionally impaired by these mutations, we took advantage of the dual regulation of *recA* transcription by both pathways. The *recA* promoter contains binding sites for both LexA and PafBC (10,17) and is therefore subject to dual regulation (Fig 1C). We introduced mutations into the SOS box (P2) or the PafBC binding sites (P1) and used these mutated promoters to drive RecA-STII expression in WT *M. smegmatis* or in strains lacking PafBC, SOS, or both (Fig 1D & E). When RecA is controlled only by SOS (P1 mutated, Fig 1D) RecA protein is induced with DNA damage (either UV or quinolone treatment) in WT cells. In cells lacking *pafBC*, RecA protein is induced by UV but not by quinolone treatment (a phenotype which will be discussed later) but RecA expression by both UV and quinolone treatment is impaired in cells lacking SOS or both pathways, indicating that the LexA-S167A strongly impairs SOS activation in vivo (Figure 1D). In contrast, when RecA is controlled only by PafBC (P2 mutated, Figure 1E), RecA protein is not induced with DNA damage in cells lacking PafBC but is inducible in WT cells as well as cells lacking SOS (Figure 1E). These experiments confirm the functional inactivation of the PafBC and SOS pathways and provide a genetic model system to dissect the relative roles of these two pathways in DNA repair and mutagenesis.

### Roles of SOS and PafBC in the transcriptional DDR across DNA damaging agents

The transcriptomic profiles of both the Δ*pafBC* and Δ*recA* strains have been assessed primarily in response to mitomycin C (MMC) in *M. smegmatis* and *M. tuberculosis* (10,12,13). However, different types of DNA damaging agents produce fundamentally different types of DNA lesions that may require unique systems for correction, the expression of which may in turn be governed by SOS or PafBC. To investigate the possibility of DNA damage specific responses for these pathways, we measured the transcriptional responses of the WT, Δ*recA*, Δ*pafBC, lexAS167A* and Δ*pafBC/lexAS167A* strains after exposure to UV or ciprofloxacin. In WT cells, UV and ciprofloxacin induced a common set of 185 genes (log_2_ fold change of >= 1.5, Supplementary Fig 1A). There were an additional 24 genes exclusively induced by cipro which were composed of genes predicted to be involved in replication, recombination, and repair. In contrast, UV induced an additional 174 genes, most of which were genes with unknown function. To deduce whether PafBC or LexA control different gene sets in response to different types of DNA damage, we focused on the subset of genes that were commonly induced by UV and ciprofloxacin (from our studies) as well as MMC from the literature in wild type cells (10).

Transcriptomic profiling of the DDR pathway mutants revealed three major DNA damage response profiles. Profile 1 consists of genes whose expression levels with DNA damage are completely dependent on the PafBC pathway, and independent of RecA, and this gene set is the largest regulon transcriptionally induced after DNA damage (Fig 2A and Table S1), consistent with prior studies (10). 35% of these genes are predicted to encode proteins involved in replication, recombination, and repair, which has been previously noted as being overrepresented in the PafBC regulon after MMC induced damage (10). Profile 2 consists of genes that have varying degrees of dependence on PafBC, SOS, or RecA (Fig 2B and Table S2). Of particular interest are the genes that are codependent on PafBC and SOS after ciprofloxacin damage, but are PafBC independent after UV (Fig 2B, red and blue boxes). This subset is of interest because they indicate DDR pathway clastogen specific responses and consist of genes with important roles in DNA damage repair and response. Although UV induces a larger number of genes compared to ciprofloxacin (Fig S1A), this pathway specific pattern is unique to ciprofloxacin. 27% of the DDR pathway ciprofloxacin specific subset have a partial or complete loss of induction in the Δ*pafBC* or *lexAS167A* strains. Of the genes with ciprofloxacin specific induction in the *lexAS167A* strain, 82% are also either fully or partially dependent on the PafBC pathway. Profile 3 consists of genes that are DNA damage inducible but have no discernible loss of expression in any of the DDR pathway mutants (Fig 2C & Table S3). A majority of these are genes with unknown functions. These results confirm prior results that PafBC controls a numerically larger number of DNA damage inducible genes irrespective of the type of DNA damage (Figures 2D & E), but also indicate that there are clastogen and pathway specific overlapping gene sets within the DNA damage response.

**Figure 2.**
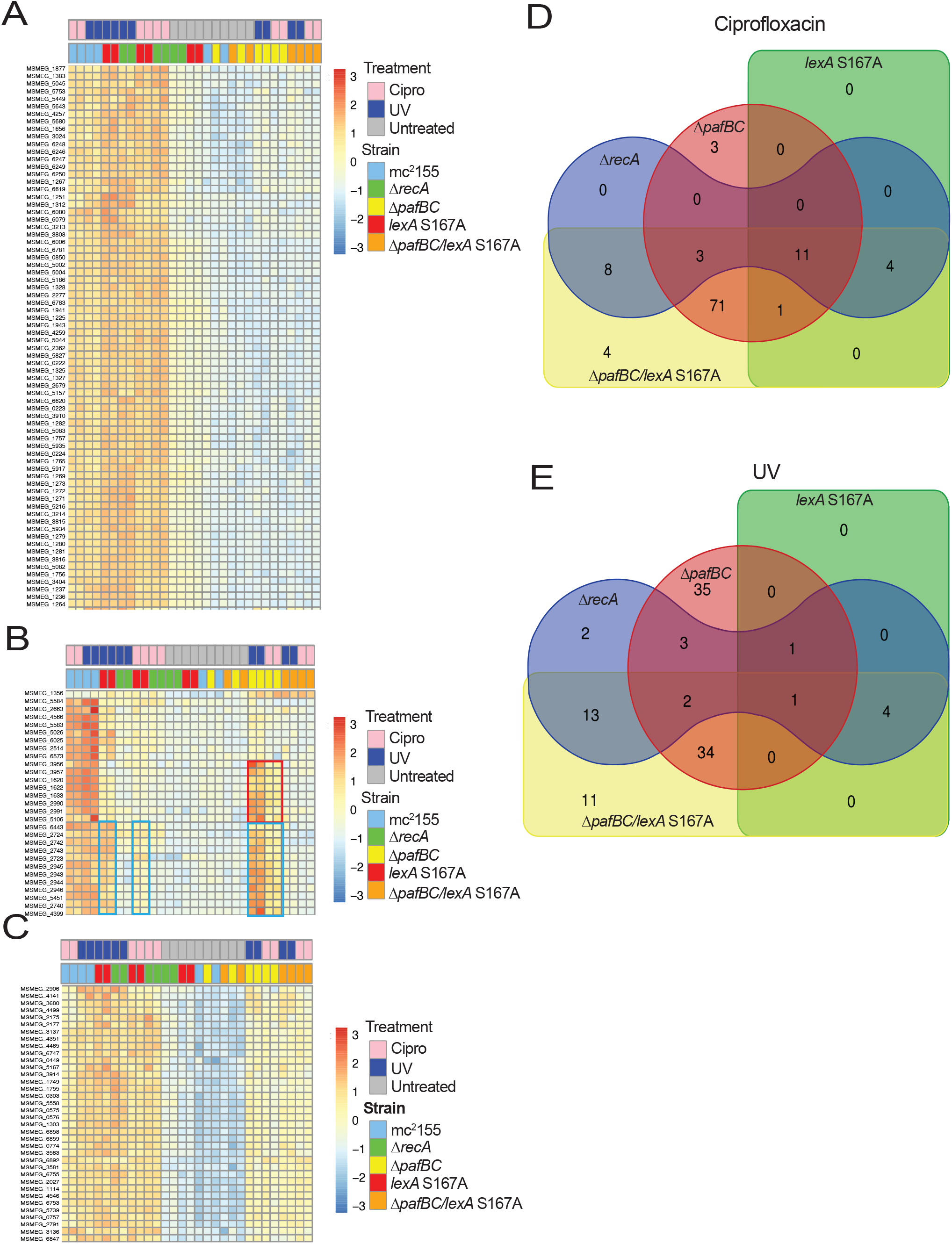
Relative contributions of PafBC or LexA to the transcriptional DNA damage response in mycobacteria. Gene expression heatmaps of genes that are significantly upregulated (log_2_ fold change >= 1.5, p-value < 0.01) in WT *M. smegmatis* mc^2^155 by ciprofloxacin (0.5μg/ml) or UV (20mJ/cm^2^) from transcriptomic profiling by RNA sequencing of mc^2^155, Δ*recA*, Δ*pafBC, lexA*-S167A and Δ*pafBC*/*lexA*-S167A strains. **(A)** The RecA independent PafBC regulon. The heat map displays genes in which DNA damage induced expression is dependent on PafBC, but preserved in LexAS167A and ΔrecA strains. The scale bar depicts normalized expression level. **(B)** Codependent and clastogen specific DDR. The red box highlights genes that are RecA/SOS/PafBC dependent with cipro, but only SOS dependent with UV. The blue boxes highlight PafBC/SOS codependent genes with cipro stress **(C)** Genes whose DNA damage dependent expression levels are independent of either DDR pathway. **(D**,**E)** Venn diagram categorizing genes for which the DNA damage induction is abolished (WT log_2_ fold change is >2.5x of the compared strain) in the indicated strain backgrounds (Δ*recA*, Δ*pafBC*, lexA-S167A and Δ*pafBC*/*lexA*-S167A) compared to mc^2^155 with ciprofloxacin (D) or UV (E) treatment. The genes represented in the Venn diagrams are the same genes represented in the heatmaps in panels A-C.

We extended these findings to *M. tuberculosis* by comparing the transcriptional response of a strain of *M. tuberculosis* H37Rv with a transposon insertion in *pafC* to both WT (H37Rv) and a *pafC*::*tn* complemented strain after exposure to both UV and ciprofloxacin. Like *M. smegmatis*, UV induced a larger number of genes in WT compared to ciprofloxacin (Fig S1B). Although the number of genes uniquely induced by ciprofloxacin alone in both *M. smegmatis* and *M. tuberculosis* WT cells being similar (Fig S1B), the only ciprofloxacin-unique gene that was induced in both mycobacterial species was *dnaQ* (Rv3711c/MSMEG_6275). Mycobacterial DnaQ is a homologue of the *E*.*coli* 3’-5’ exonuclease epsilon. To confirm the PafBC and ciprofloxacin phenotypes observed in *M. smegmatis*, we focused on the subset of genes that were induced by both UV and ciprofloxacin (Fig S1B). Although an *M. tuberculosis* LexA uncleavable strain was not available for comparison, we observed that 57% of the genes are no longer induced by either clastogen in the absence of PafBC (Table S4). We also observed a similar subset of genes that have a PafBC ciprofloxacin dependent phenotype and include *recA* (Table S4 and Fig S1C). This data confirms our observation of clastogen specific gene sets within the DNA damage response.

To confirm our RNAseq results and gain a clearer temporal picture of the roles of the SOS and PafBC pathways in the temporal transcriptional response to different DNA damaging agents, we analyzed the mRNA encoding AdnA (a PafBC dependent gene in our RNAseq dataset and in the literature (10,12)) and DnaE2 (an SOS dependent gene as defined in our RNAseq dataset and in the literature (12)) by RT-qPCR. Analyzing the expression of *adnA* after treatment with UV, ciprofloxacin or MMC confirmed the results from the transcriptomic profiling that both basal and induced *adnA* expression are dependent on the PafBC pathway, regardless of the DNA damaging agent tested, as reflected in the significantly reduced expression of *adnA* with or without DNA damage in the Δ*pafBC* and Δ*pafBC/lexA-S167A* strains (Fig 3A-C). Impairment of the SOS pathway in both the Δ*recA* and *lexAS167A* strains led to a significant increase in *adnA* expression in basal conditions (Fig 3A-C). However, induction of *adnA* in the SOS deficient strains after DNA damage was not impaired compared to wild type cells, further confirming that *adnA* is exclusively controlled by the PafBC, RecA independent pathway. The expression of *adnA* in the Δ*pafBC* strain was rescued by expressing either *M. smegmatis* or *M. tuberculosis* PafBC (Fig 3D), confirming that these phenotypes are due to loss of PafBC function. *dnaE2* is reported to be SOS dependent, a conclusion that is derived from its impaired expression in mycobacteria lacking *recA* (12). Our experiments confirm this impaired expression of *dnaE2* with DNA damage in Δ*recA* (Fig 3E-G). However, although loss of SOS in the lex*AS167A* strain impairs *dnaE2* induction to a greater degree than loss of *pafBC*, there is a significant impairment of *dnaE2* expression in Δ*pafBC*, especially with ciprofloxacin and MMC. Only in the Δ*pafBC/lexAS167A* strain does *dnaE2* expression phenocopy that observed in Δ*recA* (Fig 3E-G). The impaired expression of *dnaE2* in the Δ*pafBC* was rescued by complementation with either *M. smegmatis* or *M. tuberculosis pafBC* (Fig 3H).

**Figure 3.**
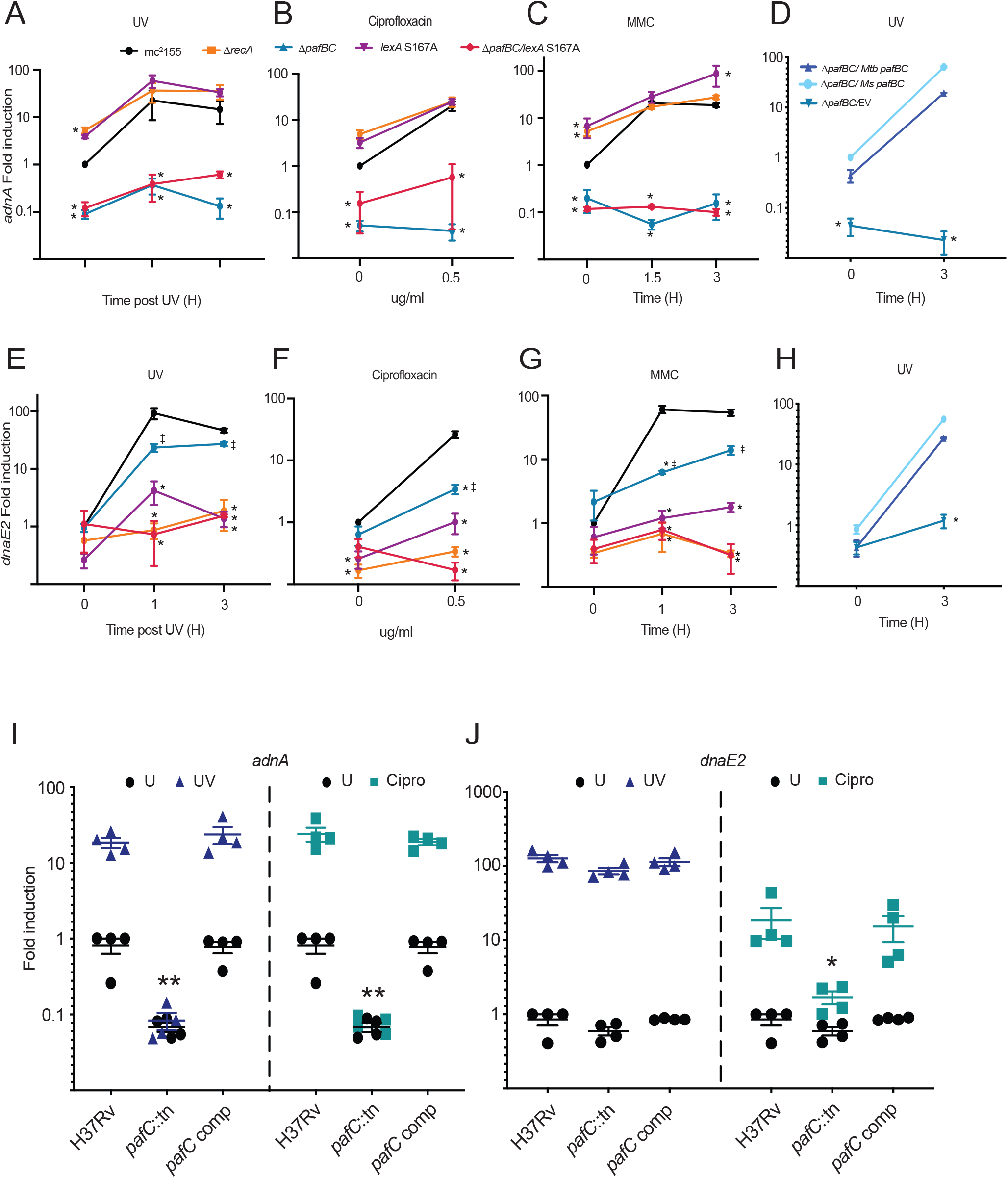
Gene and clastogen specific requirements for PafBC and LexA in the transcriptional DDR. **(A-C)** Normalized *adnA* mRNA measured by RT-qPCR in the indicated strains of *M. smegmatis* (black=wild type; orange=Δ*recA*, blue=Δ*pafBC*, purple=*lexAS167A*, and red=Δ*pafBC/ lexAS167A*) after exposure to either **(A)** UV (20mJ/cm^2^) (n= 3 biological replicates) **(B)** ciprofloxacin (0.5μg/ml) (n = 5 biological replicates) or **(C)** Mitomycin C (MMC) (80ng/ml) (n = 3 biological replicates). All values are normalized to WT untreated at 1. **(D)** Genetic complementation. Normalized *adnA* mRNA measured by RT-qPCR in *M. smegmatis* Δ*pafBC* complemented with either *M. tuberculosis pafBC, M. smegmatis pafBC* or empty vector (EV) after exposure to UV (20mJ/cm^2^) (n = 2 biological replicates). All values are normalized to *M. smegmatis* Δ*pafBC* complemented with *M. smegmatis pafBC* **(E-G)** Normalized *dnaE2* mRNA measured by RT-qPCR in the same *M. smegmatis* strains as in (A-C) after exposure to either **(E)** UV (20mJ/cm^2^) (n= 3 biological replicates) **(F)** ciprofloxacin (0.5□g/ml) (n = 5 biological replicates) or **(G)** MMC (80ng/ml) (n = 3 biological replicates). All values are normalized to WT untreated at 1 **(H)** Normalized *dnaE2* mRNA in *M. smegmatis* Δ*pafBC* strains complemented with either *M*.*tuberculosis pafBC, M*.*smegmatis pafBC* or empty vector (EV) (same strain legend as in panel D) after exposure to UV (20mJ/cm^2^) (n = 2 biological replicates). All values are normalized to *M. smegmatis* Δ*pafBC* complemented with *M*.*smegmatis pafBC* untreated at 1 **(I-J)** Normalized mRNA measured by RT-qPCR for **(I)** *adnA* or **(J)** *dnaE2* in *M. tuberculosis* H37Rv, H37Rv *pafC*::*tn, pafC*::*tn* +*pafC*. All values are normalized to WT untreated at 1. Significance is calculated as *= p < 0.05 using 2-way ANOVA compared to the WT strain (mc^2^155 or H37Rv) at a comparable timepoint/condition. * = p < 0.05 using 2-way ANOVA compared to *lexAS167A* strain at a comparable timepoint/condition. Error bars are SEM

We extended these findings to *M. tuberculosis* using the strain of *M. tuberculosis* H37Rv with a transposon insertion in *pafC*. We confirmed that *adnA* expression is reduced in basal conditions in cells lacking *pafC*, a phenotype that is complemented by the wild type gene (Fig 3I). UV or ciprofloxacin induction of *adnA* was completely abrogated in *pafC*::*tn* (Fig 3I). In contrast, although UV induced *dnaE2* expression was independent of *pafC* (Fig 3J), *dnaE2* induction with ciprofloxacin was substantially dependent on *pafC* (Fig 3J), again complemented by wild type *pafC* (Fig 3J) as is seen in *M. smegmatis*. These results mirrored the transcriptional induction of both *adnA* and *dnaE2* observed in *M. tuberculosis* by RNA sequencing (Table S4). Taken together, these data confirm the PafBC dependence of *adnA* expression, but also implicate PafBC in *dnaE2* expression under specific conditions. These data suggest that these two systems may mediate the response to different types of DNA damage and indicate that loss of *recA* has more severe effects on the DDR than specific ablation of the SOS pathway that more closely resembles loss of both SOS and PafBC. These data suggest a role for RecA in inducing the PafBC regulon under certain types of DNA damage.

### Roles of SOS and PafBC in DNA damage survival across DNA damaging agents

Although the transcriptional output controlled by PafBC or LexA is useful to understand the relative roles of these two transcription factors, the ultimate functional outcomes of the DDR may be governed by a small number of gene products or non-transcriptional mechanisms (8). To functionally characterize the relative contributions of LexA and PafBC, we measured the survival of the mutants after treatment with the same clastogens used for transcriptional profiling. As previously described (8), loss of *recA* severely sensitizes *M. smegmatis* to all forms of DNA damage, a phenotype that could partially reflect its role in SOS induction, but also other repair roles such as DSB repair and restart of stalled replication forks (18,19). Loss of LexA cleavage severely sensitized cells to UV killing, almost to the same degree as loss of *recA*, with PafBC playing a more minor role, especially at low doses (Fig 4A). Only the Δ*pafBC/lexA-S167A* strain fully phenocopied loss of *recA*. With gyrase inhibition (ciprofloxacin), the PafBC and SOS pathways contributed nearly equally, but only loss of both pathways phenocopied loss of *recA* (Fig 4B). For mitomycin, loss of LexA cleavage sensitized cells to killing more than loss of PafBC (Fig 4C). The sensitivity of the Δ*pafBC* strain was rescued by expressing *M. smegmatis* or *M. tuberculosis pafBC* (Fig 4D). These results indicate that, despite controlling a smaller number of genes, the SOS pathway plays an important role in survival after damage. In addition, the data confirms the finding from transcriptional profiling that these two pathways respond to different types of damage, with SOS having a dominant role for UV and crosslinking and PafBC playing a more important role in the response to gyrase inhibition.

**Figure 4.**
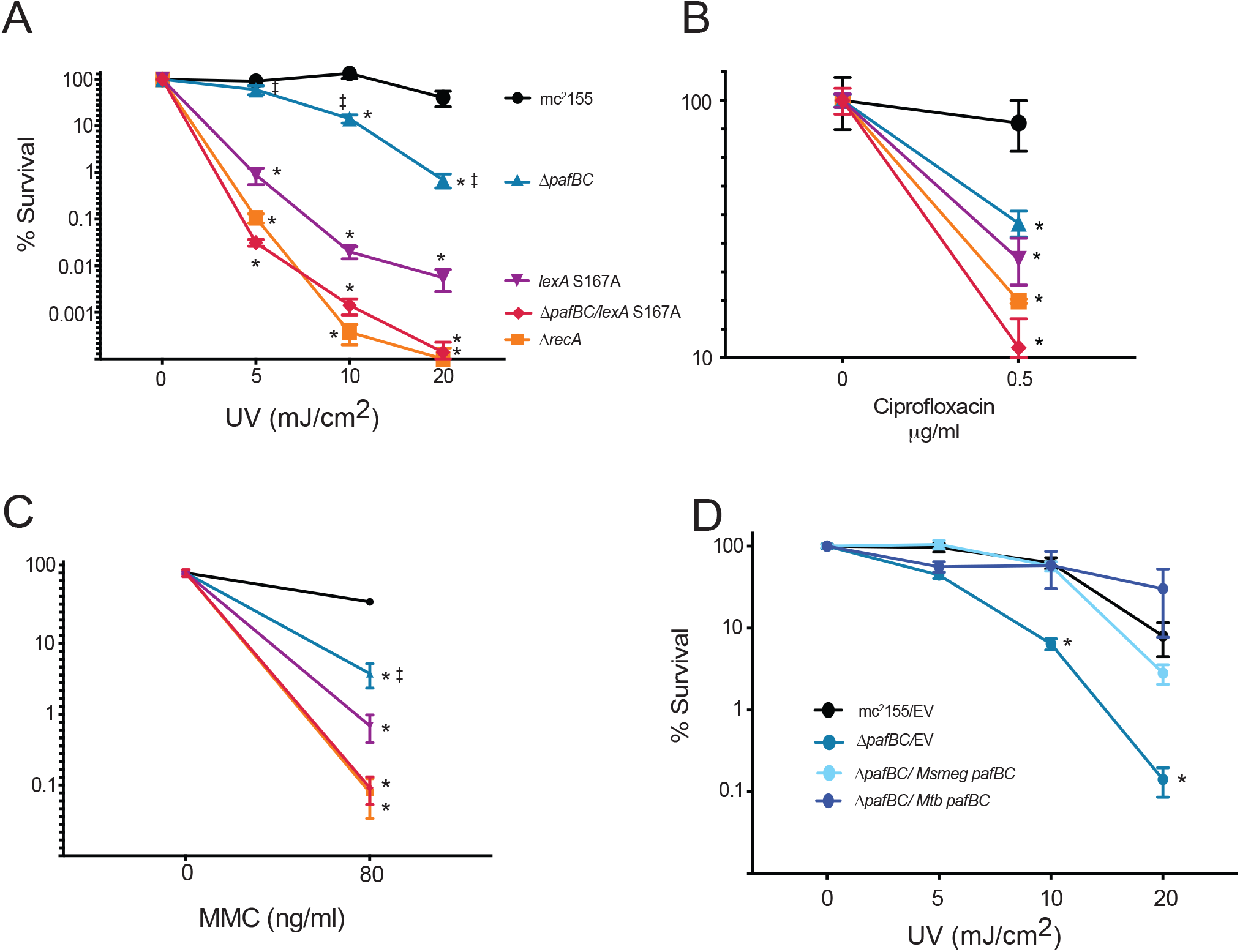
Functional outputs of the DDR are codependent on LexA and PafBC. Survival of mc^2^155 (black), Δ*recA* (orange), Δ*pafBC* (blue), *lexAS167A* (purple) and Δ*pafBC*/*lexA-S167A* (red) after exposure to **(A)** UV (0, 5, 10 or 20mJ/cm^2^) (n = 4 biological replicates) **(B)** ciprofloxacin (0.5μg/ml) (n = 4 biological replicates) or **(C)** MMC (80ng/ml) (n = 5 biological replicates) **(D)** Survival of Δ*pafBC* strains complemented with either *M. tuberculosis pafBC, M. smegmatis pafBC* or empty vector (EV) and mc^2^155 complemented with EV after exposure to UV (20mJ/cm^2^) (n = 2 biological replicates). Significance is calculated as *= p < 0.05 using 2-way ANOVA compared to the WT strain at a comparable timepoint/condition. ‡= p < 0.05 using 2-way ANOVA compared to *lexAS167A* strain at a comparable timepoint/condition. Error bars are SEM

### Roles of SOS and PafBC in adaptive mutagenesis

DnaE2 has previously been shown to play a major role in adaptive mutagenesis after UV induced DNA damage (15). As we observed that *dnaE2* expression is controlled by both SOS and PafBC when DNA damage was induced by MMC and ciprofloxacin, but not by UV, we quantitated the functional output of adaptive mutagenesis by measuring the appearance of Rifampin resistance (Rif^R^) after UV light exposure. As expected, UV strongly induces the frequency of Rif^R^ in WT cells and loss of *recA* abolishes that induction (Fig 5A) (8). Surprisingly, we found loss of either DDR pathway (Δ*pafBC* and *lexAS167A* strains) impaired mutagenesis (Fig 5A), whereas loss of both pathways eliminated mutagenesis (Fig 5A). The loss of mutagenesis in the Δ*pafBC* mutant could be rescued by complementation using *M. smegmatis pafBC* (Fig 5B). Extending these findings to *M. tuberculosis* revealed a reduction in adaptive mutagenesis in the absence of *pafC*, a phenotype that was complemented by the wild type *pafC* gene (Fig 5C). The decrement of UV-induced mutagenesis in the absence of the PafBC pathway was surprising because *dnaE2* expression was not significantly reduced after UV exposure in either *M. smegmatis* or *M. tuberculosis pafBC* single mutants (Fig 3E & 3J). Although DnaE2 activity is required for its mutagenic role in vivo (15), the mutasome also consists of the ImuA/B cassette, which are also required for mutagenesis in vivo (23). These genes are reported to be SOS regulated (12), but recent data also indicates that there is a PafBC binding site in the *imuA’* (MSMEG_1620) promoter and DNA damage induced expression of *imuA/B* was impaired in *M. smegmatis* lacking *pafBC* (10). Consistent with this prior data, our transcriptomic profiling revealed that *imuA*/*B* are strongly induced by both UV and quinolone damage and this induction is abolished in cells lacking *recA*, a finding previously interpreted to indicate SOS dependence. To clarify the regulation of *imuA/B*, we measured the expression of *imuB*, which is an operon with *imuA* (23), by RT-qPCR and found that *imuB* expression was significantly impaired by the loss of either the SOS and PafBC pathways (Figure 5D and 5E). We observed a partial impairment of *imuB* in both the *lexAS167A* and Δ*pafBC* knockout strains, with a stronger impairment conferred after ciprofloxacin induced DNA damage and only in the *pafBC*/*lexAS167A* strain did we observe complete loss of *imuB* expression that phenocopied loss of *recA* (Fig 5D). In *M. tuberculosis, imuB* expression was induced with UV exposure, and was mildly impaired in cells lacking *pafC* (Figure 5E). However, *imuB* induction after ciprofloxacin was abolished in *pafC*::*tn* cells, a phenotype that was complemented by wild type *pafC* (Fig 5E). These data indicate that both the PafBC and SOS pathways contribute to adaptative mutagenesis in mycobacteria, in part through shared regulation of the *dnaE2* cofactors *imuA/B*.

**Figure 5.**
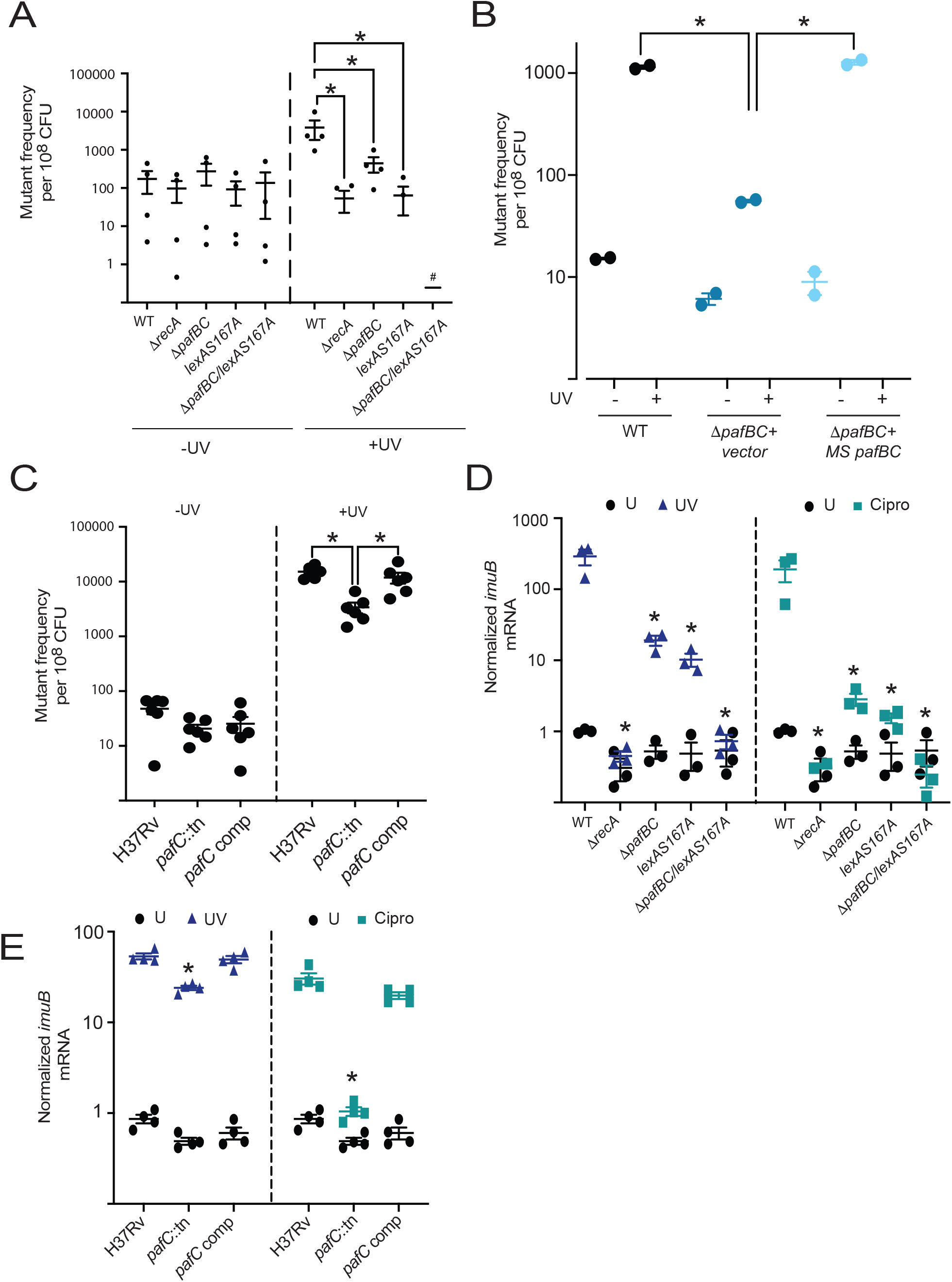
Mycobacterial mutagenesis requires both SOS and PafBC. **(A)** Frequency of rifampin resistant mutants per 10^8^ cells in *M. smegmatis* mc^2^155, Δ*recA*, Δ*pafBC, lexA-S167A* and Δ*pafBC*/*lexAS167A* with or without DNA damage (20mJ/cm^2^ UV) (n= 4 biological replicates). # indicates that no Rif^R^ colonies were recovered (**B**) Frequency of rifampin resistant mutants per 10^8^ cells in WT *M*.*smegmatis*, Δ*pafBC* or Δ*pafBC* complemented with *M*.*smegmatis pafBC* after exposure to UV (20mJ/cm^2^) (n = 2 biological replicates) (**C**) *pafBC* is required for mutagenesis in *M. tuberculosis*. Frequency of rifampin resistance per 10^8^ cells in *M. tuberculosis* strains of H37Rv, transposon insertion mutant of *pafC* (*pafC* TN) and transposon insertion mutant of *pafC* complemented with *pafC* (*pafC* comp) with or without DNA damage (20mJ/cm^2^ UV) (n= 6 biological replicates) (**D**) Normalized *imuB* mRNA measured by RT-qPCR in the indicated strains of *M. smegmatis* after exposure to either UV (20mJ/cm^2^) (n=3 biological replicates) or ciprofloxacin (0.5μg/ml) (n = 3 biological replicates). All values are normalized to WT untreated at 1 (**E**) Normalized *imuB* mRNA measured by RT-qPCR in the indicated strains of *M. tuberculosis* after exposure to either UV (20mJ/cm^2^) (n= 4 biological replicates) or ciprofloxacin (0.5□g/ml) (n = 4 biological replicates). All values are normalized to WT untreated at 1. Significance is calculated as *= p < 0.05 using 2-way ANOVA compared to WT untreated. Error bars are SEM

### Dual regulation of the RecA promoter is clastogen specific

The data presented above indicates that PafBC and SOS have both shared and independent roles in the DDR, which may reflect overlapping functionality of their regulons or dual regulation of specific gene products, as is the case for *imuA/B. recA* is one example of a gene directly regulated by both PafBC and SOS as it contains binding sites for LexA and PafBC in its promoter ((17) and Fig 1C). To further examine whether SOS and PafBC respond to distinct signals, we used RecA expression to measure the temporal output of both pathways. RecA protein expression after UV treatment was strongly induced by one hour, and although the Δ*pafBC* strain has lower baseline RecA expression without DNA damage, RecA levels reach the same level by three hours (Fig 6A). A similar pattern was observed in SOS deficient cells, indicating that both pathways control UV induced RecA expression. In contrast, ciprofloxacin treatment strongly induced RecA in wild type cells, but this induction was abolished in cells lacking PafBC, and preserved in cells lacking SOS (Fig 6B). In the case of mitomycin C, all three tested strains had similar expression levels of RecA both with and without DNA damage (Fig 6C). This ciprofloxacin specific loss of RecA induction is rescued by complementing Δ*pafBC* with either *M*.*smegmatis or M*.*tuberculosis pafBC* (Fig 6D). These findings indicate that the overlapping functions of these pathway are in part governed by the type of DNA damage. Most remarkable is the finding that quinolones appear to be specific inducers of the PafBC arm of the DDR. To confirm this finding in *M. tuberculosis*, we treated the H37Rv *pafC* transposon mutant with cipro and UV and measured RecA protein. Although UV induced RecA expression was unaffected by loss of *pafC*, quinolone induced RecA expression was completely dependent on *pafC*, consistent with the results in *M. smegmatis* (Fig 6E). These results indicate that ciprofloxacin is a specific inducer of the PafBC pathway.

**Figure 6.**
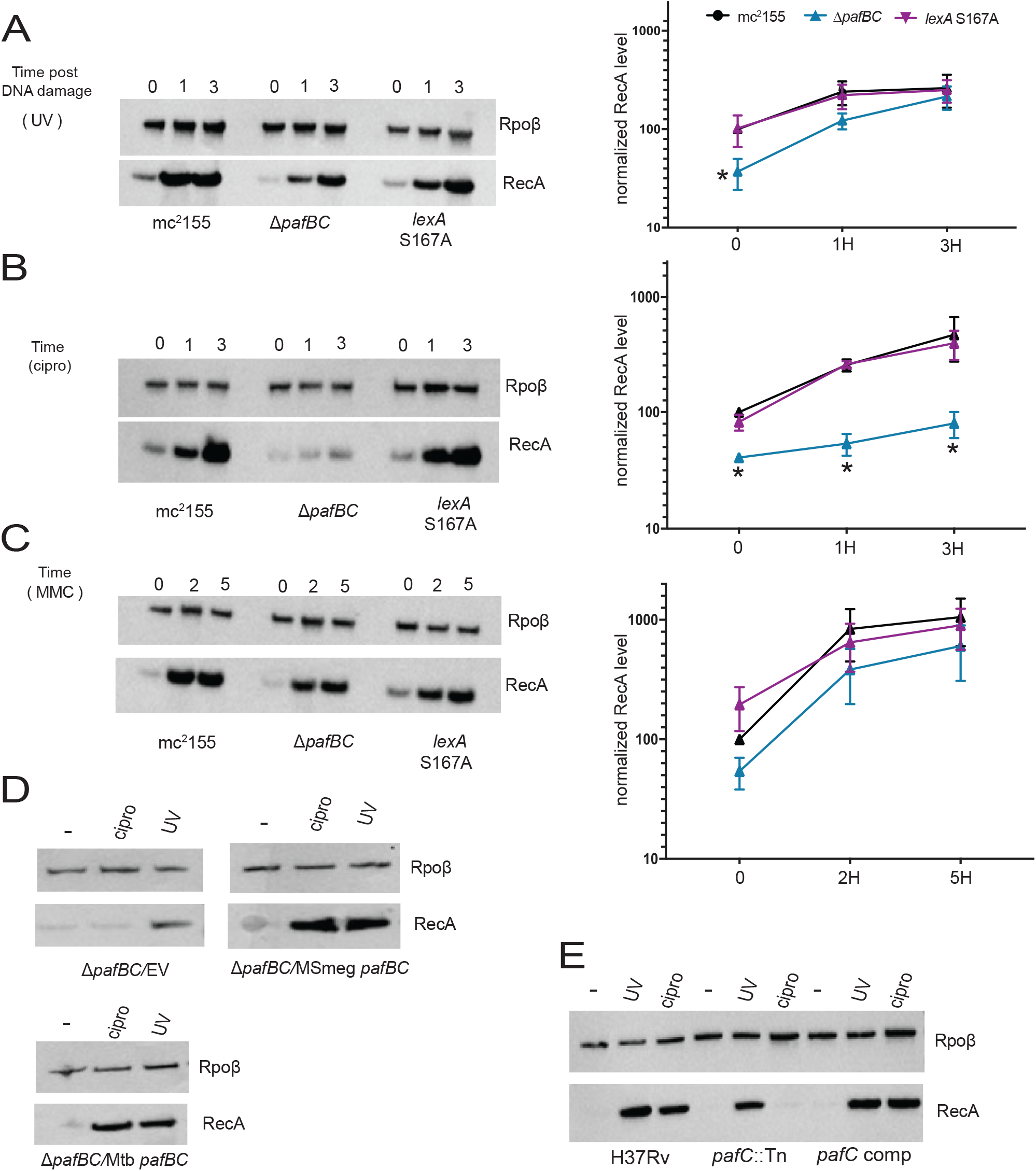
Ciprofloxacin is a selective inducer of the PafBC pathway. α-RecA western blot of mid-log phase expression of RecA (37kD) in *M. smegmatis* WT, Δ*pafBC* and *lexAS167A* after exposure to **(A)** UV (20mJ/cm^2^ with a recovery period of 1 hour or 3 hours after exposure), **(B)** ciprofloxacin (1.25μg/ml for 1 hour or 3 hours) or (C) MMC (80ng/ml for 2 hours or 5 hours). RpoB is shown as a loading control. Quantification of RecA levels normalized to RpoB levels from 3 biological replicates for each of the DNA damaging agents is shown in the right panels of (A), (B) and **(C)**. All RpoB normalized RecA levels are displayed as a percentage of WT untreated set at 100%. Significance is calculated as *= p < 0.05 using 2-way ANOVA compared to mc^2^155 at a comparable timepoint/condition. Error bars are SEM **(D)** α-RecA immunoblot of mid-log phase expression of RecA (37kD) in *M. smegmatis* Δ*pafBC* complemented with either *M. tuberculosis pafBC, M. smegmatis pafBC* or empty vector (EV) after exposure to UV (20mJ/cm^2^) or ciprofloxacin (1.25µg/ml) after 3 hours **(E)** α-RecA western blot of mid-log phase expression of RecA (37kD) in *M. tuberculosis* strains of WT H37Rv, transposon insertion mutant of *pafC* (*pafC* TN) and transposon insertion mutant of *pafC* complemented with *pafC* (*pafC* comp) after exposure to UV (20mJ/cm^2^ with a recovery period of 24 hours after exposure) or ciprofloxacin (1.25µg/ml for 24 hours). RpoB is shown as a loading control

### Damage survival and mutagenesis of SOS and PafBC are not due to impaired RecA expression

Although the data above clearly delineates overlapping and clastogen specific roles for SOS and PafBC in RecA expression, and RecA has multiple roles in DNA repair and mutagenesis, the contribution of impaired RecA expression to the functional defects observed in cells lacking SOS, *pafBC*, or both, is unknown. To examine this question, we placed RecA expression under the control of an anhydrotetracycline (ATc) inducible promoter (*irecA*) and complemented the WT, Δ*recA*, Δ*pafBC, lexAS167A* and, Δ*pafBC/lexAS167A* strains. Addition of ATc induced RecA protein accumulation at a higher level than wild type cells in the absence of DNA damage and enhanced UV induced RecA accumulation (Fig. S2A). UV treatment of the Δ*recA* strain complemented with *irecA* revealed no inducibility of RecA, but constant RecA levels with ATc (Fig S2A). i*recA* restored RecA expression in the SOS and PafBC deficient strains (Figure S2B) including the doubly deficient strain, which has nearly undetectable RecA levels.

To test whether enforced RecA expression could reverse the DNA damage sensitivity phenotypes observed with SOS and PafBC deficient cells, we measured killing with UV with and without RecA expression. Although *irecA* expression did not change the survival profile of the WT cells, it did fully rescue the severe survival defect of the Δ*recA* strain treated with UV, indicating that *irecA* encodes a functional RecA protein that is expressed at levels that can rescue complete RecA deficiency (Fig S2C). However, restoration of RecA expression had no effect on the survival of SOS, PafBC, or doubly deficient cells (Fig S2C). Similarly, *irecA* did not change the DNA damage induced transcription of *adnA* in WT cells and was unable to rescue the loss of *adnA* transcription in either the Δ*pafBC* or Δ*pafBC/lexA* S167A strains (Fig S2D). However, irecA fully rescued *dnaE2* expression in Δ*recA*, and partially rescued the transcription of *dnaE2* in the *lexAS167A* and Δ*pafBC/lexAS167A* strains (Fig S2E). These results indicate that, despite the prominent coregulation of RecA expression by PafBC and SOS, the defective RecA expression conferred by loss of these pathways does not explain their damage sensitivity.

### Replisome perturbation induces the PafBC pathway

An essential question about the PafBC pathway is its mechanism of activation. The PafBC proteins are not themselves damage inducible (10), suggesting that a signal generated by DNA damage results in PafBC activation. Whether this occurs at the level of induced DNA binding, dimerization, or some other mechanism, is unknown (24). The data above indicates that quinolones may be a specific activating signal for the PafBC pathway (Fig 6). Quinolones act by inhibiting DNA gyrase and thereby affect DNA replication and transcription, in addition to inducing protein linked double strand DNA breaks. To understand the induction of the PafBC pathway by quinolones, we tested other stresses that may impact DNA replication or transcription. Although oxidative stress induction by cumene hydroperoxide (CHP) (Fig 7A) or inhibiting DNA replication using the antimicrobial SKI-356313 (25) (Fig 7B) both induced RecA expression in WT cells, this induction was not dependent on *pafBC*. In contrast, the topoisomerase inhibitor etoposide (26) induced RecA in WT cells, and this induction was substantially dependent on PafBC, but not SOS (Fig 7C). To more precisely perturb DNA replication, we expressed the alternative lesion bypass polymerase DinB1 from an ATc inducible promoter. Mycobacterial DinB1 interacts with the DNA replication machinery beta clamp (27) and therefore may stall the replication fork when overexpressed. Consistent with this prediction, DinB1 expression strongly induced RecA (Fig 7D), but this induction was abolished when DinB1Δ356-360 was expressed, which carries a mutation in the beta-clamp interacting motif (Fig 7E). RecA induction by DinB1 overexpression was lost in Δ*pafBC but* preserved in the SOS deficient strain (Fig 7D), consistent with the hypothesis that perturbation of the replication fork is an inducer of the DDR pathway. Polymerase inactive DinB1 (*dinB1* D113A) strongly induced RecA, but this induction was no longer *pafBC* dependent (Fig 7F), suggesting that this inactive polymerase activated an alternative form of DNA damage, possibly by inhibiting access of repair factors to the fork.

**Figure 7.**
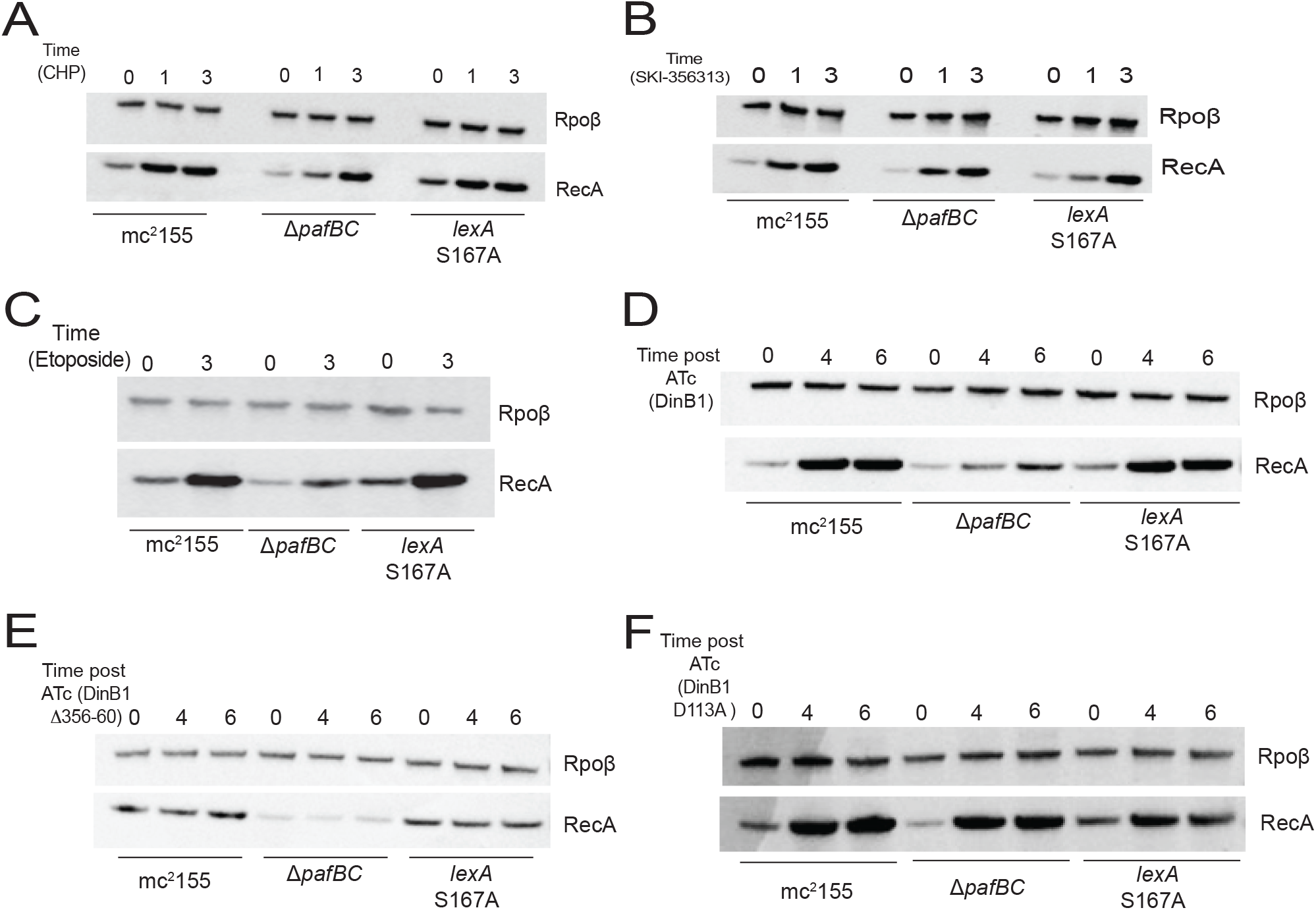
Inhibition of DNA replisome function selectively induces the PafBC pathway. α-RecA immunoblot of mid-log phase expression of RecA (37kb) in mc^2^155, Δ*pafBC* and *lexAS167A* strains after exposure to **(A**) cumene hydroperoxide (CHP, 50μM for 1 hour or 3 hours) **(B)** SKI356313 (1.9μM for 1 hour or 3 hours) **(C)** Etoposide (24uM for 3 hours **(D)** ATc induced overexpression of DinB1 for 4 hours or 6 hours **(E)** ATc induced overexpression of DinB1 missing the beta-clamp interaction domain (DinB1-Δ356-360) for 4 hours or 6 hours or **(F)** ATc induced overexpression of polymerase-dead DinB1 (*dinB1-*D113A) for 4 hours or 6 hours.

## Discussion

In this study, we have taken a comprehensive approach to examine the transcriptional and functional contributions of both the PafBC and SOS pathways in the mycobacterial DNA damage response program. Our findings reveal the requirement for both pathways to mount a full and effective response to DNA damage inflicted by various DNA damaging conditions. Our analysis of the transcriptional response of mycobacteria to distinct types of DNA damage reveal that, although the PafBC pathway controls the larger transcriptional response irrespective of the type of DNA damage, both arms of the DDR are required for survival after damage and for mutagenesis. In addition, we reveal important differences between the pathways that in part depend on the type of DNA damaging agent. Most prominently, our data indicates that quinolone antibiotics, which inhibit DNA gyrase, or replication fork perturbation, specifically induce the PafBC arm of the DDR.

Despite encoding a larger transcriptional regulon, our findings reveal that both the SOS and PafBC pathways play major roles in surviving the effects of DNA damage and in mutagenesis that results from the repair of this damage. In mycobacteria, the mutasome consists of a complex of the DnaE2 polymerase (15), along with ImuB-ImuA (23), all of which are required for the generation of rifampin resistance mutations. ImuB interacts directly with the beta-clamp of the replication apparatus and DnaE2 (23). Our results indicate that a partial loss of *imuA/B* expression in the absence of the PafBC pathway is accompanied by impaired mutagenesis in cells lacking *pafBC*. Although *imuA*/B expression is partially SOS dependent, *M. smegmatis* or *M. tuberculosis* lacking *pafBC* or *pafC* respectively display impaired *dnaE2* or *imuB* expression after DNA damage. This *pafC* dependence is particularly dramatic after gyrase inhibition, during which *dnaE2* or *imuB* expression is completely *pafBC* dependent.

This quinolone specific, *pafBC* dependent response to DNA damage has potential clinical relevance. Although the in vitro activity of quinolones against *M. tuberculosis* has long been recognized, and they have been part of second line therapy for MDR TB, recent clinical data indicates a role for the quinolone Moxifloxacin in 4-month Rifapentine regimens to treat drug sensitive TB (28). Coupled with our finding that there is a role for PafBC in supporting mutagenesis, our findings provide a molecular basis for the possibility that widespread use of quinolones for TB treatment may promote mutations that could enhance resistance to other antibiotics. In *E*.*coli*, studies have shown a requirement for the SOS pathway in inducing mutagenesis after quinolone treatment (29).

Our data provide perspective on prior efforts to understand the division of labor between the classical SOS pathway and the PafBC pathway. Based on the validated role for RecA as the LexA coprotease, and before the identification of PafBC, groundbreaking work demonstrated that the mycobacterial DDR had “RecA dependent” and “RecA independent” arms (12,30,31). The RecA dependent arm was thought to represent the SOS arm of the pathway. However, the data presented here reveals a more complex relationship between RecA and SOS. Although RecA null cells are clearly SOS null, they also are clearly more severely deficient for DNA repair and mutagenesis than cells specifically ablated for SOS. Only when SOS and PafBC are both ablated do the DNA damage phenotypes resemble loss of RecA. The activation signal for the PafBC pathway is unknown. The PafBC heterodimer is constitutively expressed and therefore one model of its activation, in part based on structural studies (24), is that a ligand generated during DNA damage binds directly to PafBC and induces a conformational change that results in DNA binding. Alternative models are possible, including sequestration of the heterodimer in basal conditions and/or induced heterodimerization. Our study does not identify the specific inducing signal for the PafBC pathway, but our data does support a model in which this signal is generated by replication fork perturbation or arrest. Given the role of RecA in replication fork restart (19), our data may suggest that in certain circumstances the activating signal for PafBC requires RecA and replication fork perturbation, a hypothesis that can be pursued by future studies.

## Acknowledgements

The authors thank Heran Darwin for providing the H37Rv *pafC* Tn and complemented strains. This work was supported by NIH grants AI 064693, P30 CA008748.

## Conflicts of interest

MG serves as an SAB member and holds equity in Vedanta Biosciences, is on the SAB of PRL-NYC, and is a consultant for Fimbrion therapeutics.

## Methods

### Bacterial strains/plasmid Constructions and Growth Conditions

*Mycobacterium smegmatis* strains are derivatives of mc^2^155 and were grown and maintained in Difco Middlebrook 7H9 media (broth) supplemented with 10% ADS (0.5% albumin, 0.085% NaCl, 0.2% dextrose) and 0.05% Tween 80 or on Difco Middlebrook 7H10 (agar) supplemented with 0.5% glycerol and 0.5% dextrose at 37°C. Gene deletions were made by homologous recombination and double negative selection (20). The point mutant of LexA was generated using the previously described oligo recombineering procedure (32). Mutant strains were confirmed by PCR using primers outside the cloned region, followed by sequencing of the amplified PCR product to confirm the strains. *Mycobacterium tuberculosis* strains are on the H37Rv background and were grown and maintained in 7H9 media (broth) or on 7H10 (agar) supplemented with 10% Oleic Acid-Albumin-Dextrose-Catalase supplement (OADC), 0.5% glycerol, and 0.01% Tyloxapol (broth only) at 37°C. For a complete strain list with relevant features, see Supplementary table 4. Plasmids utilized in this study were generated using standard molecular techniques and along with relevant oligos are listed with their features in Supplementary table 4.

### RT-qPCR

*M. smegmatis* cultures were grown to OD600 ∼ 0.5 - 0.6, collected by centrifugation at 3200G and re-suspended to OD600 = 0.6. For each treatment (0.5μg/ml Ciprofloxacin, 80ng/ml Mitomycin C or 10ml culture exposed to 20mJ/cm^2^ UV), 10ml cultures were shaken at 37 °C /150RPM at a final OD600 = 0.3 (For UV, 5ml treated culture in 5ml of fresh media) for the indicated time hours and collected for RNA preparation, lysed by bead beating 3x for 30 seconds and after a 24-hour incubation in 500ul RNAlater buffer RNA was isolated using the GeneJet RNA purification kit. 500ng of RNA (quantified on ThermoScientific Nanodrop 8000 spectrophotometer) was used to make cDNA using the Thermo Maxima First Strand cDNA synthesis kit for RT-qPCR with dsDNase. The RT-qPCR reaction was made for a Taqman assay using Thermoscientific DyNAmo Flash probe qPCR kit (10ul of mastermix, 0.1ul of each primer, 0.05ul of each probe, 5ul of cDNA sample and 4.5ul of deionized H2O per reaction) and analyzed using the Applied Biosystems 7500 Real-Time system (cycling conditions: 95°C for 7 mins, 45 cycles of 95°C for 5 seconds and 60°C for 30 seconds). Primer/probe sets for each target genes of *dnaE2, adnA* and *imuB* were combined with primer/ probe sets for *sigA* as the housekeeping gene and the analysis was done by comparing the ΔΔCT for each treated strain to WT mc^2^155 untreated control. Each cDNA sample was tested in duplicate and no RT control reactions were included in all RT-qPCR experiments to exclude spurious amplification of contaminating chromosomal DNA.

*M. tuberculosis* strains were grown to OD600 ∼ 0.5 - 0.6, collected by centrifugation at 3200G and re-suspended to OD600 = 0.6. For each treatment (0.5μg/ml Ciprofloxacin or 8ml culture exposed to ∼20mJ/cm^2^ UV), cultures with a final volume of 10ml were shaken at 37 °C /150RPM at a final OD600 = 0.3 (For UV, 5ml treated culture in 5ml of fresh media) for 24 hours, lysed in TRIzol reagent by bead beating 3 times for 45 seconds and processed using the Direct-zol Miniprep Plus kit. RNA was treated following the rigorous DNase treatment of the TURBO DNA-free kit. cDNA synthesis and RT-qPCR were similar to protocol used for *M. smegmatis* cells. Primers for RT-qPCR are in supplementary table 4.

### Ribosomal RNA depletion and Transcriptional Profiling

*M. smegmatis* and *M. tuberculosis* RNA samples for ciprofloxacin (0.5μg/ml for 3 hours (*M. smegmatis*) or 24 hours (*M. tuberculosis*)) or UV (20mJ/cm^2^ with a recovery period of 1 hour (*M*.*smegmatis*) or 24 hours (*M. tuberculosis*) after exposure) transcriptional profiling by RNA sequencing were isolated from cells grown and treated as described above. The *M. smegmatis* RNA samples (n=2) were depleted for ribosomal RNA using a biotinylated oligonucleotides-based protocol (33). The *M. tuberculosis* RNA samples(n=1) were depleted for ribosomal RNA using the Ilumina Ribozero Plus rRNA depletion kit. For both rRNA depletion methods, the efficiency of rRNA depletion was variable between samples with the percentage of reads mapped to rRNA after depletion ranging from 11% to 92%. Despite this variable rRNA depletion, all samples contained greater than 3 million non-ribosomal mapping transcripts. RNA sequencing was performed as previously reported (34).

### DNA damage assays

Strains were grown to saturation and were diluted to an OD600 = 0.02 and grown to OD600 ∼0.6, collected by centrifugation at 3200G and resuspended to an OD600 = 0.6. For UV exposure, cells were serially diluted in PBS + 0.05% tween80 onto 7H10 agar plates. Agar plates were treated with the indicated doses of UV radiation with a Stratagene UV stratalinker 1800 with 254nm UV bulbs. Plates were wrapped in foil (to prevent potential effects of photolyase) and incubated at 37°C. For treatment with ciprofloxacin (0.5μg/ml for 3 hours) and Mitomycin C (80ng/ml for 3 hours), cultures were incubated at a final OD600 = 0.3. Cultures were washed with PBS + 0.05% tween80 and then were serially diluted in PBS + 0.05% tween80 onto 7H10 agar plates

### Western Blot

*M. smegmatis* lysates were prepared from cells exposed to ciprofloxacin (1.25 μg/ml for 1 and 3 hours), UV (20mJ/cm^2^ with a recovery period of 1 and 3 hours after exposure), Mitomycin C (80ng/ml for 2 and 5 hours), Anhydrotetracycline (ATc) (50ng/ml for described time), Cumene hydroxyperoxide (CHP) (50μm for 1 hour or 3 hours), SKI356313 (1.9μm for 1 hour or 3 hours) or Etoposide (24μm for 3 hours) by bead beating 2x for 30s (5-minute rest on ice between each cycle). Protein quantities were normalized using the Protein A280 on the Nanodrop 8000 and normalized to an apparent concentration of 0.2mg/ml. Blots were blocked and probed in 5% Omniblot milk in 1XPBST (Phosphate buffered saline + 0.01% tween20). Equal loading was confirmed with commercially available Biolegend *E. coli* anti-RpoB antibody (1:10,000 dilution) as a loading control. RecA antibody, raised in rabbits against purified full-length *M. smegmatis* RecA (8), was used at a 1:5,000 dilution. For RecA-streptag II (STII), STII (GenScript, Rabbit Anti-NWSHPQFEK polyclonal antibody) was used at a 1:5,000 dilution and the blots were blocked and probed in strep wash buffer (150mM NaCl, 100mM Tris PH 7.9/8, 1mM EDTA). LexA antibody, raised in rabbits against purified full-length *M. smegmatis* LexA, was used at a 1:2,000 dilution. Blots were imaged in iBright FL1000 and quantified using the iBright analysis software. *M. tuberculosis* lysates were prepared from cells exposed to ciprofloxacin (1.25 μg/ml for 24 hours) or UV (20mJ/cm^2^ for 24 hours) by bead beating 3x for 45s. Blots were blocked and probed similarly to *M. smegmatis* samples.

### UV induced Mutagenesis

10 mL of each strain at OD_600_ = 0.6 was transferred to Omnitray single-well plates (*M. smegmatis*) or extra depth disposable petri dish (*M. tuberculosis*) and exposed to 20 mJ/cm^2^ UV radiation using a Stratagene UV stratalinker 1800. From each treated sample and its untreated control, 5 mL of culture was transferred to 5 mL of fresh media and shaken at 37 °C /150 RPM for 3 hours (*M*.*smegmatis*) or 24 hours (*M*.*tuberculosis*). From each sample, a total of 5-9ml of culture was cultured as 400μl aliquots on 7H10 agar plates containing 0.5% glycerol, 0.5% dextrose, and 100 μg/mL rifampicin and incubated at 37 °C for 72 hours (*M*.*smegmatis*) or 3 weeks (*M. tuberculosis*) to determine rifampicin-resistant CFU. Additional duplicates were taken from each sample and dilution-plated on 7H10 agar containing no antibiotic to determine viable CFU. Resistant mutants were then normalized to viable CFU for each set of samples. Graph represents average rifampicin-resistant mutants per viable CFU.

### Statistical analyses

Significance tests were performed in GraphPad Prism using a two-way analysis of variance (ANOVA) test on log-transformed values. All performed statistical tests were two-sided. All error bars represent standard error of the mean (SEM), unless specifically noted otherwise.

### Data availability Statement

RNA sequencing data has been deposited into into the SRA as BioProject# PRJNA746693.

## Figure legends

**Figure S1.**
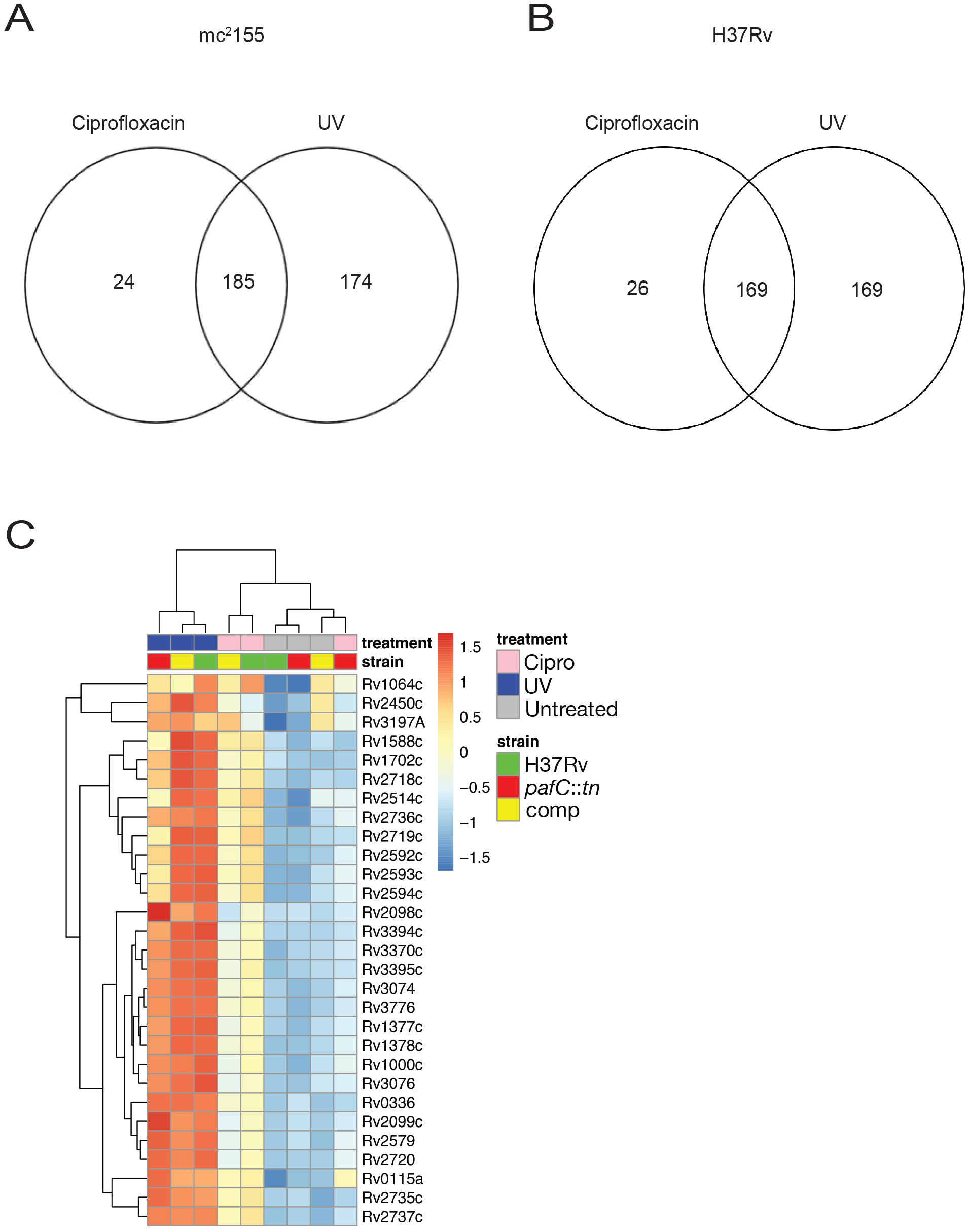
Transcriptional profiles after ciprofloxacin and UV induced DNA damage. **(A)** Venn diagram categorizing genes that are upregulated (log_2_ fold change >= 1.5) in mc^2^155 by ciprofloxacin (0.5*μ*g/ml) or UV (20mJ/cm^2^) (n = 2) **(B)** Venn diagram categorizing genes that are upregulated (log_2_ fold change >= 1.5) in H37Rv by ciprofloxacin (0.5*μ*g/ml) or UV (20mJ/cm^2^) (n = 1) **(C) The *pafC* quinolone responsive regulon**. Gene expression heatmap of genes that are induced by ciprofloxacin and are *pafC* dependent in this condition, but not after UV, from transcriptomic profiling by RNA sequencing of H37Rv, transposon insertion mutant of *pafC* (*pafC*::TN) and transposon insertion mutant of *pafC* complemented with *pafC* (comp)

**Figure S2.**
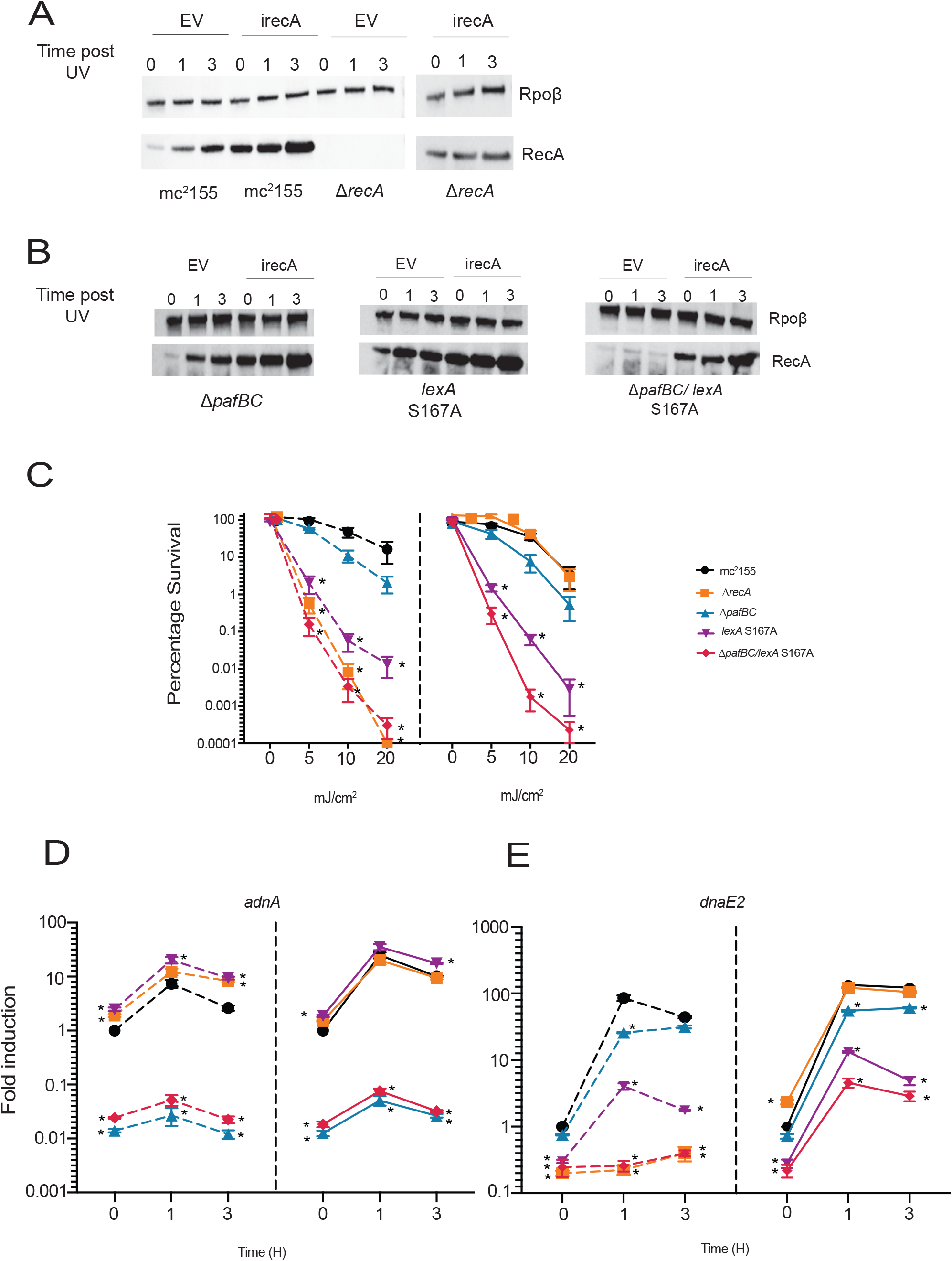
The functional impairment of the DDR that accompanies PafBC and SOS inactivation is independent of RecA. **(A)** α-RecA western blot of mid-log phase expression of RecA (37kb) in mc^2^155 and Δ*recA* strains either with an empty vector (V) or with *recA* under an ATc inducible promoter (*irecA*) after exposure to UV (20mJ/cm^2^ for 1 hour or 3 hours) and in the presence of ATc **(B)** α-RecA western blot of mid-log phase expression of RecA (37kb) in Δ*pafBC, lexA-S167A* and Δ*pafBC*/*lexA-S167A* either with an empty vector (EV) or with *recA* under an ATc inducible promoter (*irecA*) after exposure to UV (20mJ/cm^2^ for 1 hour or 3 hours) and in the presence of ATc. RpoB is used as a loading control **(C)** Percentage survival of mc^2^155, Δ*recA*, Δ*pafBC, lexA-S167A* and Δ*pafBC*/*lexA-S167A* strains with either empty vector (V: dotted lines, left panel) or with *recA* induced from an ATc inducible promoter (*irecA*: solid line, right panel) after exposure to UV (0, 5, 10 or 20mJ/cm^2^) in the presence of ATc (n = 3 biological replicates). **(D**,**E)** Normalized mRNA levels in *M. smegmatis* mc^2^155, Δ*recA*, Δ*pafBC, lexA-S167A* and Δ*pafBC*/*lexA-S167A* strains with either empty vector (EV: dotted line) or with *recA* under an ATc inducible promoter (*irecA*: solid line) after exposure to UV (20mJ/cm^2^) measured by RT-qPCR for (D) *adnA* (n= 3 biological replicates) or (E) *dnaE2* (n= 3 biological replicates). Significance is calculated as *= p < 0.05 using 2-way ANOVA compared to mc^2^155 at a comparable timepoint/condition. All error bars are SEM

